# Somatic Programmed DNA Elimination is widespread in free-living Rhabditidae nematodes

**DOI:** 10.1101/2025.08.21.671558

**Authors:** Caroline Launay, Eva Wenger, Brice Letcher, Marie Delattre

## Abstract

All cells of a multicellular organism usually share an identical genome, faithfully transmitted through successive divisions. Yet, a number of animal species deviate from this dogma, as parts of their DNA are systematically eliminated in all their somatic nuclei, in a process called Programmed DNA Elimination (PDE). PDE leads to the unexpected reorganisation of the genome at every generation in all somatic cells but its molecular mechanism, evolutionary origins, and functional significance remain unknown. This lack of understanding partially stems from limitations in genetically tractable model species.

PDE can target an entire chromosome, or involve chromosome fragmentation followed by selective fragment retention and elimination, raising further questions on genome stability, genome integrity and mechanisms of DNA repair. PDE by chromosome fragmentation has been described in parasitic nematodes in the family Ascarididae, copepods in the genus Cyclops and unicellular ciliates. More recently, PDE has been discovered in three non-parasitic, lab-tractable nematode species from the Rhabditidae family, opening new perspectives.

In this study, we used cytological approaches to screen 25 new Rhabditidae species for PDE. We found evidence of PDE in 17 species. Our work reveals that PDE is present in 12 out of 17 tested genera, demonstrating its widespread presence in Rhabditidae nematodes, with the notable exception of *C. elegans*. Genetic tools have already been established for some species. This work provides a collection of lab-tractable species that can be used to test many aspects of somatic Programmed DNA Elimination by chromosome fragmentation in animals.

## Introduction

Some species systematically eliminate portions of their genome in their soma, in a process called Programmed DNA Elimination (PDE) [1,2]. PDE can target an entire chromosome, as in songbirds or lamprey for instance [3,4]. PDE can also happen by the fragmentation and partial destruction of chromosomes, as spotted for the first time in 1887 in the parasitic nematode *Parascaris univalens* [5]. PDE by chromosome fragmentation was later identified across the phylum of ciliate protists, Ciliophora [6]. Ciliates are unicellular organisms with two nuclei: a transcriptionally inactive micronucleus (MIC) transmitted to descendants after a sexual phase, and a transcriptionally active macronucleus (MAC), copied from the MIC and amplified. During MAC development, ciliates undergo extensive PDE, often involving both excision of internal sequences, healed by non-homologous end-joining, as well as chromosome fragmentation, as found in ascarid parasitic nematodes. PDE is pervasive in ciliates. Its study in a few species has led to seminal discoveries on the role of small RNAs in genome rearrangement [6]. It also revealed that internal eliminated sequences are remnants of transposable elements, excised by a domesticated transposase [7–9]. This, together with many internal eliminated sequences being also transposon-derived, has led a widely accepted scenario that excision of internal sequences emerged as a selfish, TE-driven process in ciliates [10]. In striking contrast, PDE through chromosome fragmentation remains poorly understood in ciliates, and its molecular dissection in animals has been largely precluded because ascarids are obligate parasites and not experimentally tractable. Moreover, the functional importance of somatic PDE is also unknown because it has not been possible to abolish PDE in any animal species so far.

Recently, PDE was identified in two nematode genera belonging to the Rhabditidae family, in a different sub-order than the one including ascarids. In *Oscheius tipulae*, telomere-to-telomere genome reconstruction revealed that 0.6 % of the genome is eliminated at chromosome ends [11,12]. Almost concomitantly, and by serendipity, we discovered substantial PDE in two other Rhabditidae species - although distant from *O. tipulae* - using cytology initially: *Mesorhabditis belari* and *Mesorhabditis spiculigera*. These two species eliminate approximately 30% and 20% of their genome, respectively, at all chromosome ends but also in large internal chromosomal blocks, leading to an increased somatic karyotype, as shown in Ascarids [13].

We could easily follow chromosome fragmentation in *Mesorhabditis* embryos, across the successive embryonic divisions. Indeed, in a large range of nematode species, including ascarids and Rhabditidae, the embryonic cell lineage is highly reproducible and conserved. Moreover, the germline precursor cell is easily identified: it is the smallest and most posterior cell; it does not divide until late in embryogenesis, or even after hatching in some species; and it harbours highly compacted chromatin [14]. Somatic cells also divide in a relatively conserved order, allowing for easy identification of cell types from one species to the other, by analogy with what has been well described in *C. elegans* [14]. In *Mesorhabditis* embryos, as previously described in ascarids, we found that PDE starts with chromosome fragmentation in all five somatic precursor cells, successively and independently, as they enter mitosis. This is followed by fragment exclusion during mitosis and DNA destruction later in development. We also found in *Mesorhabditis*, as shown for ascarids or *O. tipulae*, that telomeres of the germline inherited chromosomes are systematically eliminated in the soma, and that new telomeres are assembled on the new extremities of all somatic chromosomes [12,13,15].

The finding that free-living nematode species undergo PDE following the same mechanism as in ascarids demonstrates that PDE is not a parasitic oddity in nematodes and raises the possibility that PDE, rather than having evolved independently in two distantly related clades, could be more widespread than anticipated in nematodes.

In this study, we asked whether PDE is widespread in family Rhabditidae, which originated approximately 200 million years ago [16]. Using simple DNA staining, we reveal that 17 out of 25 newly tested species show evidence of PDE. This corresponds to 12 out of 17 genera in which at least one species undergoes PDE. This work reveals that PDE is widespread in Rhabditidae, with the notable exception of *C. elegans*, and has gone unnoticed for more than a century of nematode research.

## Results

### Identification of Programmed DNA Elimination by DNA staining

In ascarids and *Mesorhabditis* species, DNA elimination is immediately visible in early embryos because chromosomes are fragmented into large blocks [5,13,17]. Prior to elimination, no fragments are visible and somatic nuclei harbour the same number of chromosomes as germ cells. During elimination, chromosome fragments can be seen excluded from the spindle during mitosis or are visible in the cytoplasm in interphasic cells. After elimination, DNA fragments are eventually destroyed after a few cell cycles and old embryos do not show cytoplasmic DNA fragments [13,15]. When fragments internal to chromosomes are also eliminated, as in ascarids or *Mesorhabditis*, the chromosome number is increased in somatic nuclei compared to germline nuclei. Overall, it is possible to detect PDE using simple DNA staining on embryos at different time points during their development.

In order to detect PDE, we performed DNA staining on embryos from 25 new species. These species belong to 17 different genera in the Rhabditidae family, which include previously studied *Oscheius* and *Mesorhabditis* genera. We unambiguously identified cytoplasmic DNA fragments in 17 Rhabditidae new species spanning 12 genera (Fig. 1, Supp. Fig. 1).

**Figure 1:**
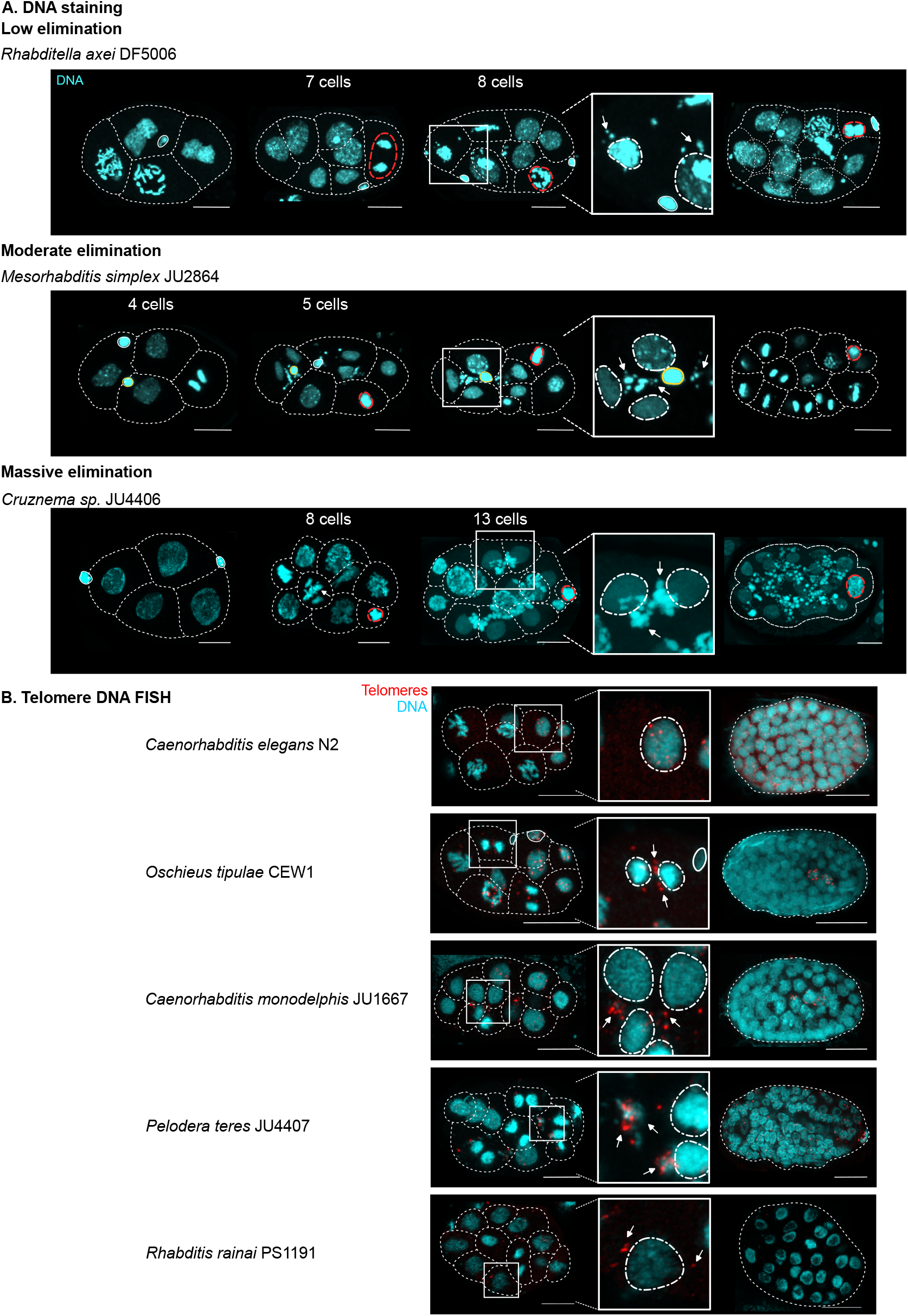
Somatic chromosome fragmentation and telomere elimination is found in several Rhabditidae species. **A**. Embryos from different species and at various stages of development (from the youngest on the left to the oldest on the right) were fixed and stained. DNA is shown in blue. The contours of the cells are shown in dotted white lines. The DNA of the germline precursor cell is shown in red. Polar bodies, when visible, are circled with a solid white line. For the pseudogamous species *Mesorhabditis simplex*, the paternal DNA stay highly condensed after fertilization [33] and is shown circled with a solid yellow line. White arrows point towards cytoplasmic fragments that become visible from the 5-cell stage in *Mesorhabditis simplex* and later in *Cruznema sp*. and *Rhabditella axei*. The name of the strain for each species is shown in parentheses. A zoomed-in view of the image of each embryo in the third column is shown in the inset on the right. In the inset, the nuclei are encircled in white and the DNA fragments excluded from the nuclei are indicated by an arrow. **B**. Embryos from different species are shown and at various stages of development, after DNA FISH treatment against telomeric repeats. On the left, embryos are undergoing chromosome fragmentation in a least one of their somatic cells. Enlargement of one cell is shown in the inset. DNA is in blue and telomeres are in red. In the inset, the nuclei are encircled in white and the telomeres excluded from the nuclei are indicated by an arrow. Older embryos, after chromosome fragmentation are shown on the right. In *Caenorhabditis elegans*, all cells in young and old embryos maintain their telomeres, whereas in species undergoing PDE, somatic telomeres do not reappear (*Oscheius tipulae, Caenorhabditis monodelphis*) or reappear progressively (*Rhabditis rainai* and *Pelodera teres*).

We reconstructed the dynamics of fragment production in all species (Fig.1 and Supp. Fig. 1). We found that fragments were initially absent in 1, 2 and 4-cell stage embryos. Fragments then appeared rapidly (possibly during a single mitotic division) and, for any given strain, reproducibly in time. After a few cell divisions, fragments were no longer detectable suggesting they had been physically destroyed. Importantly, fragments were not detected in the germ cell primordium (Fig.1 and Supp. Fig. 1). For some species we could confirm that in cells that have generated cytoplasmic fragments, the number of chromosomes in their nuclei was higher than the number of chromosomes found in early embryos or in germ cells (Supp. Fig. 1). For instance, we counted 2n=12 chromosomes in the germline of *Pelodera teres* JU4407, and found more than 26 small fragments in somatic cells from old embryos (Supp Fig. 1). This is compatible with a scenario where for each chromosome, at least one internal chromosomal fragment has been eliminated, as shown in ascarids or *Mesorhabditis* [13,15]. We ruled out that our observed fragments corresponded to intracellular bacteria, as cytoplasmic DNA fragments appeared and disappeared specifically in somatic cells, and not in the germline primordium.

We found large variation in the amount of eliminated DNA (Fig. 1 and Supp. Fig. 1). In some cases, we found very large and numerous fragments, as in *Mesorhabditis simplex*, or more dramatically even in *Cruznema sp*. JU4406 (Fig. 1). For other species, the amount of eliminated DNA seems more limited as fragments are small. This is notably the case in *Rhabditella axei* (Fig.1), *Oscheius dolichura* and *Caenorhabditis monodelphis* (Supp. Fig. 1). With our DNA staining method, we did not detect any fragments in *O. tipulae*, most likely because it eliminates 0.6% of its genome only (Supp. Fig. 1) [11,12]. This suggested that we may have underestimated the number of eliminating species using this method.

We next asked if as found in ascarids, *Mesorhabditis belari* and *Oscheius tipulae*, germline telomeres are also eliminated from somatic cells in our newly identified species. To this end, we performed DNA-FISH against the conserved telomeric repeat TTAGGC in several species (Fig. 1). We confirmed germline telomere elimination in a few species only, as the FISH signal was often too noisy to unambiguously conclude (see Material and Method). However, germline telomere elimination could be confirmed in *Pelodera teres, Rhabditis rainai, Oscheius tipulae, Auanema freiburgensis* and *Caenorhabditis monodelphis*. As a control, we showed that germline telomeres are maintained in *C. elegans* somatic nuclei, as expected from a species that does not undergo PDE (Fig. 1). We also found maintenance of germline telomeres in *Pelodera strongyloides*, a species for which we did not detect cytoplasmic DNA fragments, strongly suggesting that this species also does not undergo PDE (Supp. Fig. 1). In some species, following DNA elimination the telomere signal did not reappear in somatic nuclei, including in *Oscheius tipulae* (Fig. 1), and in *Mesorhabditis belari*, confirming our previous finding [13]. This is consistent with the detection, from sequencing reads, of shorter neo-telomeres at the new somatic chromosome ends than in the germline in *Oscheius tipulae* [11]. In contrast, the telomeric signal progressively reappears in the somatic nuclei of late embryos in *Pelodera teres* and *Rhabditis rainai*, suggesting that somatic telomeres are longer in these species than in *Mesorhabditis* species. This will need to be confirmed using genomics.

Altogether, our results reveal that at least 17 out of these 25 newly studied Rhabditidae species undergo PDE, that they most likely all eliminate germline telomeres, and in some cases additionally eliminate internal parts of chromosomes as well.

### Evolution of PDE within Rhabditidae

To identify phylogenetic patterns of PDE presence/absence, we generated a new phylogeny containing the 28 species we studied here (*M. belari, M. spiculigera, O. tipulae* and 25 new species), using 18S and 28S ribosomal DNA (rDNA) sequencing data (Fig. 2). Our phylogeny is consistent with previous publications [18], except on one occasion that we discuss below.

**Figure 2:**
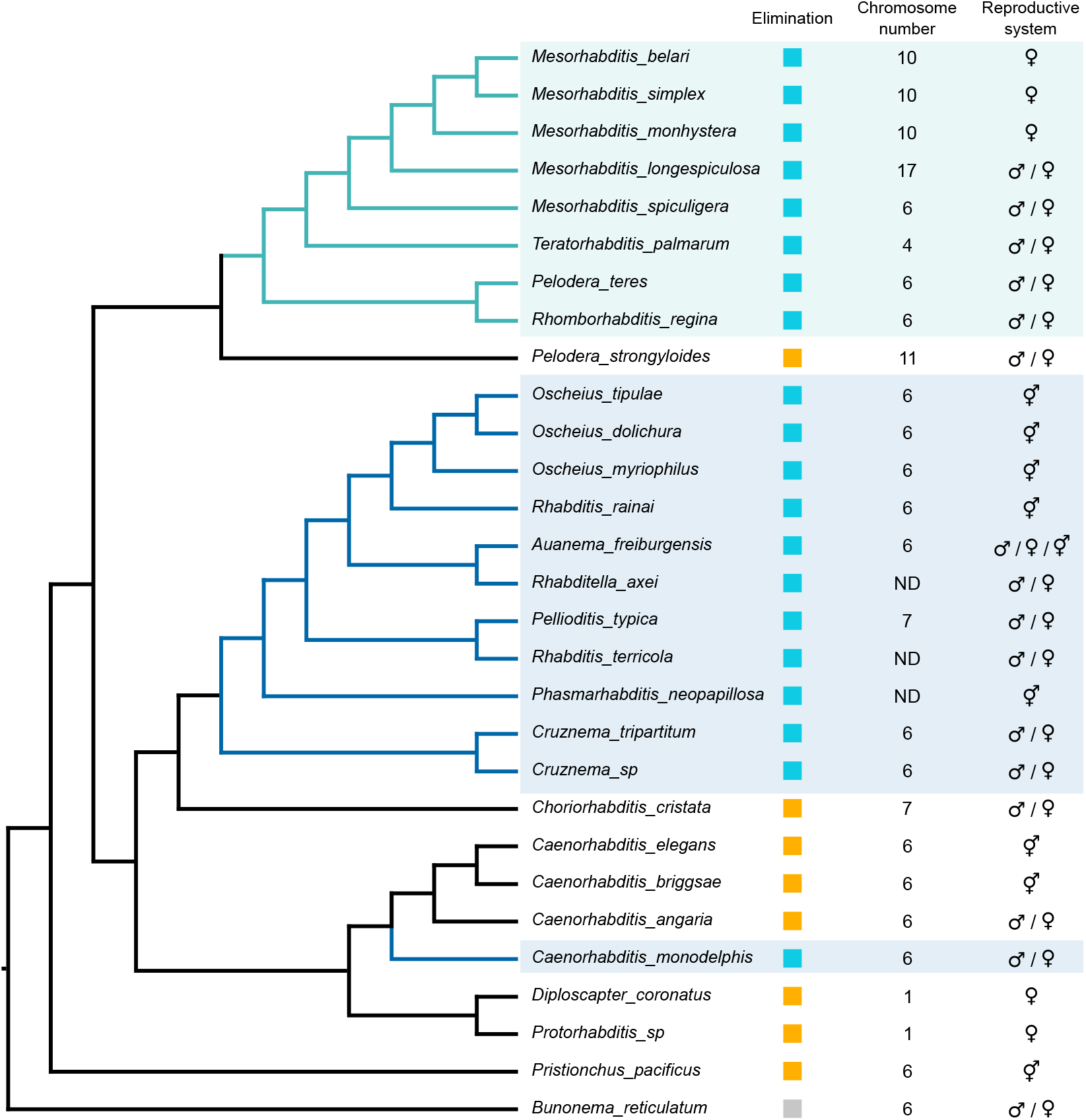
Phylogenetic distribution of species undergoing Programmed DNA Elimination in family Rhabditidae. Cladogram of species analysed in this study, based on the 18S and 28S rDNA sequencing. Species in which we detected PDE are shown in blue and others in orange. PDE has not been scored in *Bunonema reticulatum*, which serves here as an outgroup for the phylogeny only. Species undergoing PDE are grouped in two main monophyletic groups. We also found that *Caenorhabditis monodelphis* undergoes PDE whereas other *Caenorhabditis* species do not. The reproductive modes are shown on the right (asexual, male/female, hermaphrodite and three sexes), as well as chromosome numbers (also visible on Supp. Fig. 1). *Pelodera strongyloides* is not monophyletic with *Pelodera teres*, (see discussion in the main text).

We found that PDE occurs in two large monophyletic groups, both with strong bootstrap support (Fig. 2 and Supp. Fig. 2). The first clade encompasses the *Mesorhabditis* genus plus three other genera: *Pelodera, Teratorhabditis* and *Rhomborhabditis*. All five analysed *Mesorhabditis* species undergo PDE. However, we found that *Pelodera strongyloides* does not eliminate whereas *Pelodera teres* does. We note that the phylogenetic placement of *Pelodera strongyloides* is ambiguous in our tree, due to highly divergent 18S and 28S sequences (see Material and Methods). Previous publications also reported weak branch support for this specific species [18,19]. It thus remains to be determined whether *P. teres* and *P. strongyloides* truly belong to the same genus, and where exactly *P. strongyloides* branches. At this stage, we do not know if PDE was present in the common ancestor of this clade and subsequently lost in the lineage leading to *P. strongyloides* (Fig. 2), or whether it emerged after the divergence from *P. strongyloides*.

The second large monophyletic clade with PDE encompasses the *Oscheius* genus plus seven other genera (Fig. 2). All ten studied species from this clade undergo PDE, with striking variability in the amount of eliminated DNA, ranging from very little in *O. tipulae* to massively in *Cruznema sp*. JU4406 (Fig. 1). During our survey, we also found that DF5015, another putative strain of *Cruznema tripartitum* displayed much less eliminated DNA than JU4406 (Supp. Fig. 1). While this suggested strong intra-species variability, crosses between JU4406 and DF5015 produced dead embryos only, likely suggesting they are in fact different species (see Materials and Methods). Overall, our results suggest that the extent of PDE can vary over a relatively short evolutionary time scale, i.e. at least within the same genus (Fig. 2).

Outside of these two clades with many tested eliminating species, we also detected PDE in a third clade with one eliminating species, *Caenorhabditis monodelphis*. In this clade, we found PDE is absent in *C. elegans, C. briggsae* and *C. angaria*, demonstrating again variability at the genus level. In a recent study, deeper investigation within this genus confirmed using genomics that PDE occurs in *C. monodelphis*, as found here, and also identified PDE in two other basal *Caenorhabditis* species, *C. auriculariae* and *C. parvicauda*, with cytological confirmation in *C. auriculariae* [20]. That study found an absence of PDE in all other *Caenorhabditis* species - including the three we tested here - leading to the conclusion that PDE was subsequently lost in the genus *Caenorhabditis* [20].

Finally, we did not detect PDE in four other species, each belonging to a different genus: *Diploscapter sp., Protorhabditis sp., Choriorhabditis cristata* and *Pristionchus pacificus*. The position of *Pristionchus pacificus* is consistent with previous publications placing it basal to all other species studied here [16,18]. However, the position of *Choriorhabditis cristata* is inconsistent with one previously published phylogeny [18],that also placed it basally, while here *C. cristata* falls between the *Caenorhabditis* genus and the clade containing *Oscheius*. Future whole genome-based phylogenies will enable definitively resolving this inconsistency. .

Irrespective of the placement of *C. cristata*, our results suggest that PDE has been lost and/or gained at least three times independently within the family Rhabditidae (Fig. 2).

### Variation in the timing of PDE

In ascarids, chromosomes start to be fragmented in ABa and ABp cells at the 4-cell stage [5,15,17]. We found the same early fragmentation starting in ABa and ABp (the two largest and most anterior cells) in *Teratorhabditis palmarum* and also in all *Mesorhabditis* species, although embryos already had five blastomeres when ABa and ABp started dividing in *Mesorhabditis* [13] (Supp. Fig. 1). However, DNA fragments are visible only when embryos reach ∼9 cells for *Pelodera teres* and *Rhomborhabditis regina*. This delay of approximately one cell cycle for the onset of DNA fragmentation was also observed in all species of the second large clade and in *C. monodelphis. Mesorhabditis, Pelodera* and *Rhomborhabditis* are closer phylogenetically than either is to *C. monodelphis* (Fig. 2). Thus, the timing of PDE initiation varies between species, but with no obvious ties to the topology of the phylogeny.

### PDE occurs regardless of reproductive systems or chromosome number

Last, we analysed PDE in the context of the reproductive system or the chromosome number of the species, two other parameters that were known [14,21] or that we collected during our analysis (see Materials and Methods and Supp. Fig. 1). Various reproductive systems are found in Rhabditidae nematodes: regular sexual reproduction, hermaphroditism, a combination of the two with the presence of three sexes [22], strict parthenogenesis (full asexuality) [23] or autopseudogamy (pseudo-sexuality) [24]. We found no obvious link between PDE presence/absence and reproductive systems (Fig. 2). For instance, some sexual species undergo PDE(e.g. *Pelodera teres)*, while others do not (e.g. *Pelodera strongyloides* and *Caenorhabditis angaria)*. Similarly, PDE is also found in species that reproduce through all reproductive modes: hermaphroditism (e. g. *Oscheius* species), species with three sexes such as *Auanema freiburgensis*, or through autopseudogamy such as *Mesorhabditis belari* and *Mesorhabditis simplex*, but is not found in other hermaphrodites (e.g. *C. elegans*) or asexuals (e.g. *Diploscapter*).

Finally, we found no correlation between the occurrence of PDE and the number of germline chromosomes (Supp. Fig. 1 and Fig. 2). PDE can occur in species with a large number of germline chromosomes, such as *Mesorhabditis longespiculosa* (n=17) or a small number, such as *Teratorhabditis palmarum* (n=4). Conversely, PDE is found neither in *Diploscapter coronatus*, a species with a single pair of germline chromosomes, nor *Pelodera strongyloides* that has 11. Many species in Rhabditidae have six chromosomes, some of which undergo PDE, such as *Caenorhabditis monodelphis* or *Oscheius dolichura* and others that do not, such as *Caenorhabditis elegans* (Fig. 2).

### Discussion

Programmed DNA Elimination (PDE) was first described in 1887 in a parasitic nematode [5] and we had recently uncovered the same process in free-living *Mesorhabditis* species that eliminate up to 30% of their genome in the soma [13]. In this study, we revealed PDE in 17 other free-living species (out of 25 newly tested species). This corresponds to 12 out of 17 genera in which at least one species undergoes PDE. We have discovered that PDE is widespread in Rhabditidae. To confirm this finding, more species in this family can be tested, beyond our collection. Our work also reveals that this process was missed by early pioneers of nematode research, even before *C. elegans*-an outsider when considering PDE-was chosen as a model system [25].

Within Rhabditidae, we found three distinct clades in which PDE is found in at least one species. Given our current sampling, it is not possible to conclude whether PDE has emerged independently three times within this family, was present in the common ancestor with independent losses, or a combination of both. In particular, we have found that the amount of eliminated DNA can vary substantially within a genus, calling for extending observations to more species per genus. Nevertheless, if we consider PDE in ascarids, it is tempting to speculate that PDE is an ancient process in nematodes and has been lost several times, including in the well-studied species *C. elegans* [20]. An interesting parallel is the unicellular ciliates, an extremely broad phylum in which most studied representatives so far have been found to undergo PDE, except in genus *Loxodes* [26]; this includes species in genera *Blepharisma, Euplotes, Oxytricha, Tetrahymena* and *Paramecium* [6,27]. Here too, the timing and amount of eliminated DNA vary widely. To what extent PDE is ancestral in ciliates, and/or evolved repeatedly and convergently, remains to be fully elucidated.

In the case of nematodes, exploration of more species - in particular, outside the Rhabditidae family - will be required to further explore the history of PDE. Identification of the mechanisms of the molecular effectors of PDE, none of which are known so far in nematodes, in all these branches will also shed light on its evolutionary trajectory.

Interestingly, elimination of the germline telomeres is a feature shared by ascarids, *O. tipulae, Mesorhabditis*. Indeed *O. tipulae* which eliminates very little DNA and does not fragment its chromosomes internally, eliminates all its germline chromosome telomeres. We confirmed this feature in four other species and it has also been confirmed for *Caenorhabditis parvicauda* and *Caenorhabditis auriculariae* [20] To date, no species has been found to eliminate exclusively internal portions while retaining chromosome ends, indicating that telomere elimination may represent a conserved, ancestral feature.

We found variation in the developmental timing at which chromosome fragments appear in the cytoplasm. One possibility is that fragmentation begins at the same stage in all species, but that fragments are subsequently processed or sorted at different times, preventing their cytoplasmic detection. Alternatively, this variation may reflect species-specific differences in the timing at which the PDE machinery is transcribed or becomes active.

Within our collection, *C. elegans* appears as one of a minority of species that does not undergo PDE. Perhaps our understanding of PDE would have been different if the scientific community had picked another Rhabditidae species as a model species, in the mid-20th century [28]. Nevertheless, our work now reveals that PDE is found in many free-living species, all of which are potentially lab-tractable. Genetic tools are already available in some of them, including *Oscheius tipulae* [12,29] and *Auanema* [30,31], and we have also established CRISPR and RNAi in some *Mesorhabditis and Pelodera* species (unpublished). This at last opens avenues for the mechanistic exploration of the PDE in animals.

We originally discovered PDE in *Mesorhabditis* by serendipity when analysing chromosome inheritance in young embryos [13]. Our study calls for a systematic search for signs of PDE by chromosome fragmentation, using cytological approaches as described here, when embryos are easy to stain, or using bioinformatic approaches in any animal species (for example using the tool delfies [32]), because PDE could be more widespread than anticipated even outside nematodes.

## Funding

This work has been supported by a grant from the ANR-22-CE12-0027 to MD, and by a grant from the Ecole Normale Supérieure de Lyon.

## Authors contributions

MD designed the experiments. BL established the phylogeny. CL and EW performed the experimental work and built the figures. BL and MD wrote the manuscript. We acknowledge the contribution of the imaging plateform PLATIM from SFR Biosciences Lyon. We also thank Garance Fayon, Clara Bourgeay, Heloise Brun and Basile Bergeron for technical help.

## Competing interests

All the authors declare having no competing interests.

## Material and methods

### Species collection

*Strains* are maintained at 20°C on NGM plates seeded with *E. coli* OP50, following *C. elegans* protocols, as described in [33]. Strains are all listed in Supplementary Figure 1. Strains were obtained from the CGC stock center, or were kind gifts from Marie-Anne Félix or from David Fitch.

### Genetic Crosses

DF5015 and JU4406 were both matching *Cruznema tripartitum* on public databases, based on the 18S sequencing. Because they revealed contrasting amounts of eliminated DNA, we wondered whether they belong to the same species. To test this, we crossed 3 virgin females from one strain with one male of the other strain. In both cross directions, females produced dead embryos only. From this result, we concluded that DF5015 and JU4406 belong to two distinct species, though they are probably closely related as males and females can still mate.

### DNA staining and DNA FISH and immunno-staining

Samples were fixed with methanol and the freeze-cracking method, as described in [13]. We fixed embryos to analyse DNA fragments or count chromosomes. We also fixed dissected female gonads in order to count the number of chromosomes in meiotic cells. At this stage, chromosomes are compact and paired as homologs, facilitating counting. After washing in PBS, the slides were counter-stained with Hoechst (0.5 µg/ml). For DNA-FISH, we used the following fluorescent oligo probes at 500 nM (ordered at Sigma) to detect telomeres: 5’ [TAM]TTAGGCTTAGGCTTAGGCTTAGGC 3’. In many species, the telomeric signal was found inside nuclei, as expected, but also outside nuclei, which we interpreted as noise. This prevented us from concluding on the presence/absence of telomeres. Images were taken on a Zeiss LSM-800 confocal.

### Phylogeny based on 18S and 28S sequences

#### Sequencing and sequence extraction

To build a phylogeny of the 28 species we studied in family Rhabditidae, we amplified the DNA sequences of 18S and 28S ribosomal RNA by PCR using universal primers, as described in [18].

In some cases, PCR amplification or sequencing did not work or gave truncated sequences. In these cases, we turned to public databases to obtain sequences from the same species that we studied by cytology. In these cases, we used the SILVA database of ribosomal gene sequences, release 138.2, [34], downloading all SSU (18S) and LSU (28S) sequences for taxon Chromadorea (taxon ID: 119089), a superset of Rhabditidae. For any given species, when multiple hits were available in SILVA, we used the hit referenced in [18] in which the authors built a phylogeny of many of the same species of Rhabditidae as studied here, also using 18S and 28S sequences (plus the large subunit of RNA Polymerase II where available). These hits can be found in Table S1 of Kiontke *et al*. (2007)[18].

We summarise the origin of each sequence for each species we studied in table 1 below. All sequences are available on Zenodo: https://doi.org/10.5281/zenodo.18890442

**Table 1.** Origin and strain of the 18S and 28S rDNA sequences used to build our Rhabditidae phylogeny. For each of our 28 species (rows), the origin columns show whether we used our sequencing, where successful (‘This study’), or if publicly-available data was used, in which case we provide the NCBI Nucleotide accession. The sequenced strains, of our study, or of the publicly-available data, are also provided. ‘-’ indicates we did not sequence and no public data was available - this was only the case for 28S sequences.

| Species | 18S origin | 18S strain | 28S origin | 28S strain |
| --- | --- | --- | --- | --- |
| *Mesorhabditis belari* | This study | JU2817 | This study | JU2817 |
| *Mesorhabditis simplex* | This study | JU2864 | - | - |
| *Mesorhabditis monhystera* | This study | JU2855 | - | - |
| *Mesorhabditis longespiculosa* | This study | DF5017 | This study | DF5017 |
| *Mesorhabditis spiculigera* | This study | AF72 |  |  |
| *Teratorhabditis palmarum* | U13937.1 | DF5019 | This study | DF5019 |
| *Pelodera strongyloides* | U13932.1 | DF5022 | - | - |
| *Pelodera teres* | AF083002.1 | EM437 | This study | JU4407 |
| *Rhomborhabditis regina* | This study | DF5012 | This study | DF5012 |
| *Oscheius tipulae* | KP756939.1 | CEW1 | EU195969.1 | CEW1 |
| *Oscheius dolichura* | EU196010 | JU460 | EU195971.1 | JU460 |
| *Oscheius myriophilus* | U81588.1 | DF5020 | AY602176.1 | DF5020 |
| *Rhabditis rainai* | AF083008.1 | PS1191 | EU195966.1 | DF5091 |
| *Auanema freiburgensis* | KY680647.1 | SB372 | KY680646.1 | SB372 |
| *Rhabditella axei* | U13934.1 | DF5006 | AY602177.1 | DF5006 |
| *Rhabditis terricola* | This study | JU4408 | EF417152.1 | NK3 |
| *Pellioditis typica* | U13933.1 | DF5026 | EU195962.1 | DF5025 |
| *Phasmarhabditis neopapillosa* | This study | JU1104 | - | - |
| *Cruznema tripartitum* | U73449.1 | DF5015 | EU195974.1 | SB202 |
| *Cruznema sp.* | This study | JU4406 |  |  |
| *Choriorhabditis cristata* | EU196013.1 | SB300 | EU195976.1 | SB300 |
| *Caenorhabditis elegans* | This study | N2 | This study | N2 |
| *Caenorhabditis briggsae* | U13929.1 | PB102 | AY604481.1 | PB800 |
| *Caenorhabditis angaria* | JN636068.1 | PS1010 | AY604482.1 | PS1010 |
| *Caenorhabditis monodelphis* | JN636070.1 | SB341 | AY602170.1 | SB341 |
| *Diploscapter coronatus* | This study | JU4418 | BBTA01000078.4838 | PDL0010 |
| *Protorhabditis sp.* | This study | JB122 | EU195958.1 | JB122 |
| *Pristionchus pacificus* | AF083010.1 | PS312 | EU195982.1 | PS312 |
| *Bunonema reticulatum* | EU196017.1 | PDL1002 | This study | PDL1002 |

We note from Table 1 that, in a number of cases, the publicly-available sequences we used did not come from the same strain that we studied by cytology (see Supp. Figure 1), and/or the public sequences we used were not from the same strains between 18S and 28S. However, we checked our resulting phylogeny against previously-built phylogenies of Rhabditidae, and found our final phylogeny to be consistent with previous knowledge, except in one instance that is discussed above; validating the use of different strains when the strain we imaged was not available. In addition, when we used our own sequencing, we consistently found that our sequencing matched almost identically with publicly-available sequences of the same species (where available).

#### Phylogeny building

For each set of gene sequences (18S and 28S), we next produced multiple sequence alignments using mafft v7.490 [35] (‘--adjustdirection --reorder’), and trimmed the alignment with clipkit v2.3.0 [36] to remove aligned positions with more than 80% gaps (-m gappy -g 0.8). We then built a phylogeny using IQ-TREE v2.3.6 [37] with auto-detection of the best-fitting substitution model (here, GTR+F+I+R3 for both the 18S tree and the 28S tree) and 1000 bootstrap replicates (-B 1000). We then used weighted ASTRAL v1.23 [38](--treeweights) to combine the two gene trees into a species tree, using a hybrid weighting scheme (--mode 1) that accounts for both bootstrap support and branch lengths to resolve gene tree discordance, and using 16 rounds of placement and subsampling (-R). We also gave twice as much weight to the 28S tree compared to the 18S tree, as the 18S tree placed the *Diploscapter*/*Protorhabditis* group incorrectly relative to the *Caenorhabditis* group (Supp. Fig. 2).

Phylogenetic trees were visualised and rendered with FigTree (https://github.com/rambaut/figtree/).

The raw Newick-formatted tree files, and code to generate the trees from the input sequences, are available on Zenodo: https://doi.org/10.5281/zenodo.18890442

## Figure legend

**Supplementary Figure 1: Cytological results for all 28 analysed species**

Embryos from all species analysed in this study are shown. DNA is in blue. The contours of the cells are shown in dotted white lines. Polar bodies, when visible, are circled with a solid white line. For the pseudogamous species *M. belari, M. simplex, M. monhystera* [33] *and Protorhabditis sp. (*unpublished), the paternal DNA stay highly condensed after fertilization [33] and is shown circled with a solid yellow line. White arrows point towards cytoplasmic fragments. On the left, embryos at the 4 or 5-cell stage are shown. On the right, are shown embryos at the time of chromosome fragmentation for species undergoing PDE (shown with a star) or at equivalent stage for species in which we never detected cytoplasmic fragments. Images used to quantify chromosome number are shown as well. In most cases, we analysed meiotic cells during diakinesis of prometaphase. At this stage, chromosomes are paired and we therefore count a haploid number of chromosomes. For some species, it was possible to count chromosome number in 1 or 2 cell embryos during prometaphase. In this case, we counted a diploid number. Of note, for *Diploscapter coronatus* the 2 DNA bodies observed in diakinesis correspond to the two unpaired chromosome. This species as a single chromosome which does not pair during meiosis. For *Oscheius tipulae, Auanema sp. VSL2220* and *Caenorhabditis monodelphis*, the images come from DNAFISH stainings, revealing the telomeres in red. This allows validating elimination in these species which eliminate very little DNA. In *Teratorhabditis palmarum*, the number of chromosomes, some being very small, is more than 2n=8 in mitotic cells, revealing that chromosome fragmentation starts at the 4-cell stage (embryos on the left).

**Supplementary Figure 2: Phylogeny of studied species using 18S and 28S sequences**

Phylogenies of species analysed in this study, based on the 18S rDNA sequencing or 28S rDNA sequencing

## Notes

https://doi.org/10.5281/zenodo.18890442

### Competing Interest Statement

The authors have declared no competing interest.

### Summary of Updates

The phylogeny of species has been updated and consequently, Figure 2 and Supp Figure 2 have been modified. Figure 1 and Supp Figure 1, with accompagnying caption, have been slightly modified . The main text and abstract have been slightly modified throughout to improve clarity, and better justify and explain the phylogenetic relation ship between species. The main results have not been modified and the take home message has not changed

## References

1. Wang J, Davis RE. Programmed DNA elimination in multicellular organisms. Curr Opin Genet Dev. 2014;27: 26–34. doi:10.1016/j.gde.2014.03.012

2. Dedukh D, Krasikova A. Delete and survive: strategies of programmed genetic material elimination in eukaryotes. Biol Rev Camb Philos Soc. 2022;97: 195–216. doi:10.1111/brv.12796

3. Smith JJ, Timoshevskaya N, Ye C, Holt C, Keinath MC, Parker HJ, et al. The sea lamprey germline genome provides insights into programmed genome rearrangement and vertebrate evolution. Nat Genet. 2018;50: 270–277. doi:10.1038/s41588-017-0036-1

4. Kinsella CM, Ruiz-Ruano FJ, Dion-Côté A-M, Charles AJ, Gossmann TI, Cabrero J, et al. Programmed DNA elimination of germline development genes in songbirds. Nat Commun. 2019;10: 5468. doi:10.1038/s41467-019-13427-4

5. Boveri T. Ueber Differenzierung der Zellkerne wahrend der Furchung des Eies von Ascaris megalocephala. Anat Anz. 1887;2: 688–693.

6. Bétermier M, Klobutcher LA, Orias E. Programmed chromosome fragmentation in ciliated protozoa: multiple means to chromosome ends. Microbiol Mol Biol Rev. 2023;87: e0018422. doi:10.1128/mmbr.00184-22

7. Baudry C, Malinsky S, Restituito M, Kapusta A, Rosa S, Meyer E, et al. PiggyMac, a domesticated piggyBac transposase involved in programmed genome rearrangements in the ciliate Paramecium tetraurelia. Genes Dev. 2009;23: 2478–2483. doi:10.1101/gad.547309

8. Bischerour J, Bhullar S, Denby Wilkes C, Régnier V, Mathy N, Dubois E, et al. Six domesticated PiggyBac transposases together carry out programmed DNA elimination in Paramecium. Elife. 2018;7: e37927. doi:10.7554/eLife.37927

9. Nowacki M, Higgins BP, Maquilan GM, Swart EC, Doak TG, Landweber LF. A functional role for transposases in a large eukaryotic genome. Science. 2009;324: 935–938. doi:10.1126/science.1170023

10. Arnaiz O, Mathy N, Baudry C, Malinsky S, Aury J-M, Denby Wilkes C, et al. The Paramecium germline genome provides a niche for intragenic parasitic DNA: evolutionary dynamics of internal eliminated sequences. PLoS Genet. 2012;8: e1002984. doi:10.1371/journal.pgen.1002984

11. Gonzalez de la Rosa PM, Thomson M, Trivedi U, Tracey A, Tandonnet S, Blaxter M. A telomere-to-telomere assembly of Oscheius tipulae and the evolution of rhabditid nematode chromosomes. G3 (Bethesda). 2021;11: jkaa020. doi:10.1093/g3journal/jkaa020

12. Dockendorff TC, Estrem B, Reed J, Simmons JR, Zadegan SB, Zagoskin MV, et al. The nematode Oscheius tipulae as a genetic model for programmed DNA elimination. Curr Biol. 2022;32: 5083-5098.e6. doi:10.1016/j.cub.2022.10.043

13. Rey C, Launay C, Wenger E, Delattre M. Programmed DNA elimination in Mesorhabditis nematodes. Curr Biol. 2023;33: 3711-3721.e5. doi:10.1016/j.cub.2023.07.058

14. Haag ES, Fitch DHA, Delattre M. From “the Worm” to “the Worms” and Back Again: The Evolutionary Developmental Biology of Nematodes. Genetics. 2018;210: 397–433. doi:10.1534/genetics.118.300243

15. Wang J, Veronezi GMB, Kang Y, Zagoskin M, O’Toole ET, Davis RE. Comprehensive Chromosome End Remodeling during Programmed DNA Elimination. Curr Biol. 2020;30: 3397-3413.e4. doi:10.1016/j.cub.2020.06.058

16. Picao-Osorio J, Bouleau C, Gonzalez De La Rosa PM, Stevens L, Fekonja N, Blaxter M, et al. Evolution of developmental bias explains divergent patterns of phenotypic evolution in two nematode clades. Proc Natl Acad Sci USA. 2025;122: e2507529122. doi:10.1073/pnas.2507529122

17. Estrem B, Wang J. Programmed DNA elimination in the parasitic nematode Ascaris. PLoS Pathog. 2023;19: e1011087. doi:10.1371/journal.ppat.1011087

18. Kiontke K, Barrière A, Kolotuev I, Podbilewicz B, Sommer R, Fitch DHA, et al. Trends, Stasis, and Drift in the Evolution of Nematode Vulva Development. Current Biology. 2007;17: 1925–1937. doi:10.1016/j.cub.2007.10.061

19. Sudhaus W. Phylogenetic systematisation and catalogue of paraphyletic “Rhabditidae” (Secernentea, Nematoda). Journal of Nematode Morphology and Systematics. 2011;14: 113–178.

20. Stevens L, Sun S, Haruta N, Xiao L, Uwatoko N, Kieninger M, et al. Programmed DNA elimination was present in the last common ancestor of Caenorhabditis nematodes. bioRxiv; 2025. p. 2025.10.23.681605. doi:10.1101/2025.10.23.681605

21. Kiontke K. The phylogenetic relationships of Caenorhabditis and other rhabditids. WormBook. 2005 [cited 30 Jan 2016]. doi:10.1895/wormbook.1.11.1

22. Kanzaki N, Kiontke K, Tanaka R, Hirooka Y, Schwarz A, Müller-Reichert T, et al. Description of two three-gendered nematode species in the new genus Auanema (Rhabditina) that are models for reproductive mode evolution. Scientific Reports. 2017;7. doi:10.1038/s41598-017-09871-1

23. Fradin H, Kiontke K, Zegar C, Gutwein M, Lucas J, Kovtun M, et al. Genome Architecture and Evolution of a Unichromosomal Asexual Nematode. Curr Biol. 2017;27: 2928-2939.e6. doi:10.1016/j.cub.2017.08.038

24. Grosmaire M, Launay C, Siegwald M, Brugière T, Estrada-Virrueta L, Berger D, et al. Males as somatic investment in a parthenogenetic nematode. Science. 2019;363: 1210–1213. doi:10.1126/science.aau0099

25. Nigon V. Modalités de la reproduction et déterminisme du sexe chez quelques nématodes libres. 1949;11: 1–105.

26. Seah BKB, Singh A, Vetter DE, Emmerich C, Peters M, Soltys V, et al. Nuclear dualism without extensive DNA elimination in the ciliate Loxodes magnus. Proc Natl Acad Sci U S A. 2024;121: e2400503121. doi:10.1073/pnas.2400503121

27. Singh M, Seah BKB, Emmerich C, Singh A, Woehle C, Huettel B, et al. Origins of genome-editing excisases as illuminated by the somatic genome of the ciliate Blepharisma. Proc Natl Acad Sci U S A. 2023;120: e2213887120. doi:10.1073/pnas.2213887120

28. Félix M-A. RNA interference in nematodes and the chance that favored Sydney Brenner. J Biol. 2008;7: 34. doi:10.1186/jbiol97

29. Felix MA. Oscheius tipulae. WormBook. 2006; 1–8. doi:10.1895/wormbook.1.119.1

30. Yamashita T, Pires-daSilva A, Oomura S, Kusano T, Haruta N, Hasumi M, et al. Microparticle Bombardment as a Method for Transgenesis in Auanema and Tokorhabditis. MicroPubl Biol. 2025;2025. doi:10.17912/micropub.biology.001585

31. Adams S, Pathak P, Shao H, Lok JB, Pires-daSilva A. Liposome-based transfection enhances RNAi and CRISPR-mediated mutagenesis in non-model nematode systems. Sci Rep. 2019;9: 483. doi:10.1038/s41598-018-37036-1

32. Letcher B. delfies: a Python package for the detection of DNA breakpoints with neo-telomere addition. Journal of Open Source Software. 2025;10: 7385. doi:10.21105/joss.07385

33. Launay C, Félix M-A, Dieng J, Delattre M. Diversification and hybrid incompatibility in auto-pseudogamous species of Mesorhabditis nematodes. BMC Evol Biol. 2020;20: 105. doi:10.1186/s12862-020-01665-w

34. Silva. [cited 21 Aug 2025]. Available: https://www.arb-silva.de/

35. Katoh K, Misawa K, Kuma K, Miyata T. MAFFT: a novel method for rapid multiple sequence alignment based on fast Fourier transform. Nucleic Acids Res. 2002;30: 3059–3066. doi:10.1093/nar/gkf436

36. Steenwyk JL, Iii TJB, Li Y, Shen X-X, Rokas A. ClipKIT: A multiple sequence alignment trimming software for accurate phylogenomic inference. PLOS Biology. 2020;18: e3001007. doi:10.1371/journal.pbio.3001007

37. Nguyen L-T, Schmidt HA, von Haeseler A, Minh BQ. IQ-TREE: A Fast and Effective Stochastic Algorithm for Estimating Maximum-Likelihood Phylogenies. Mol Biol Evol. 2015;32: 268–274. doi:10.1093/molbev/msu300

38. Zhang C, Mirarab S. Weighting by Gene Tree Uncertainty Improves Accuracy of Quartet-based Species Trees. Mol Biol Evol. 2022;39: msac215. doi:10.1093/molbev/msac215

