## Supplementary figures and images for "Somatic Programmed DNA Elimination is widespread in free-living Rhabditidae nematodes"

### Supp Figure 1

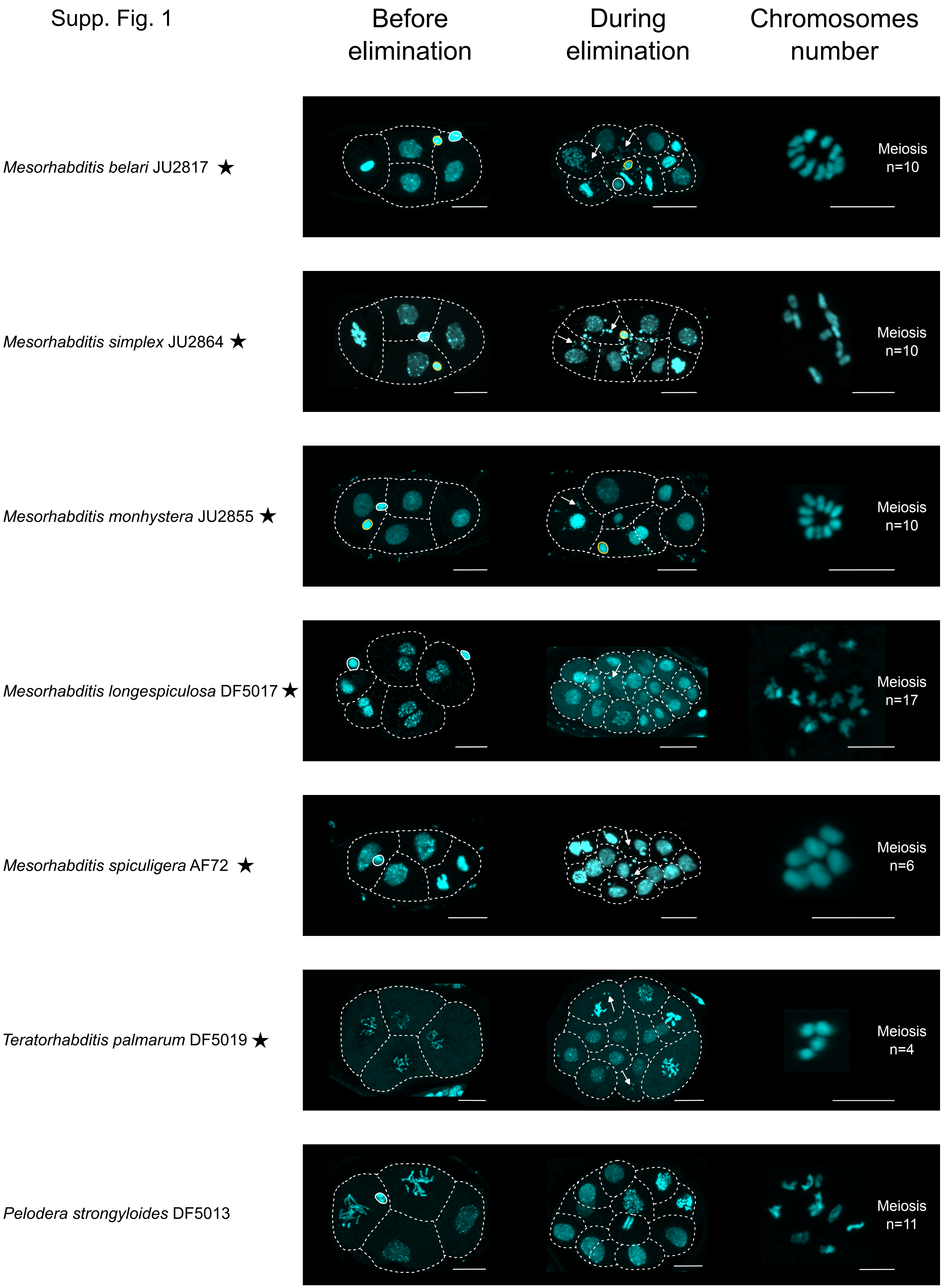

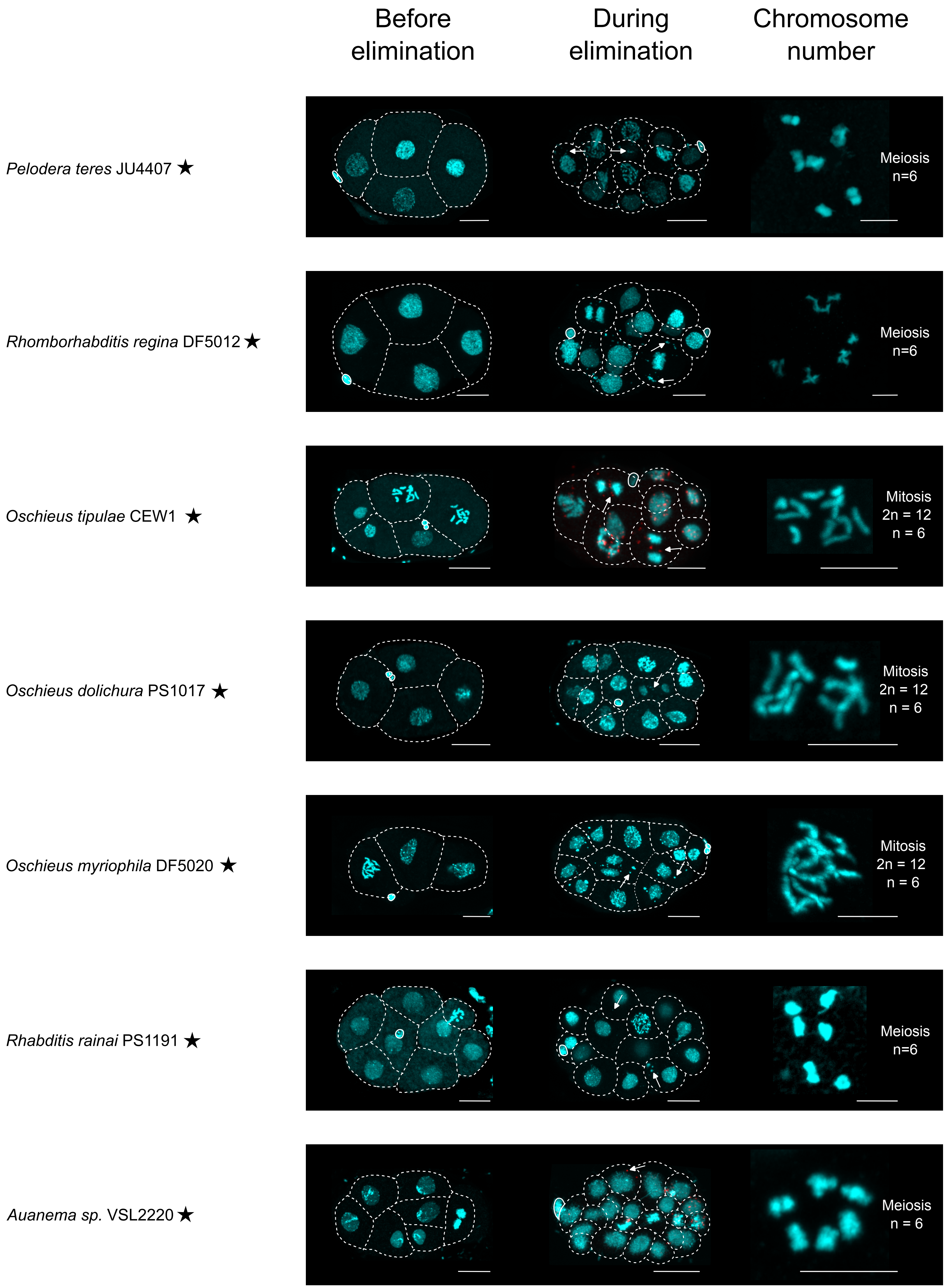

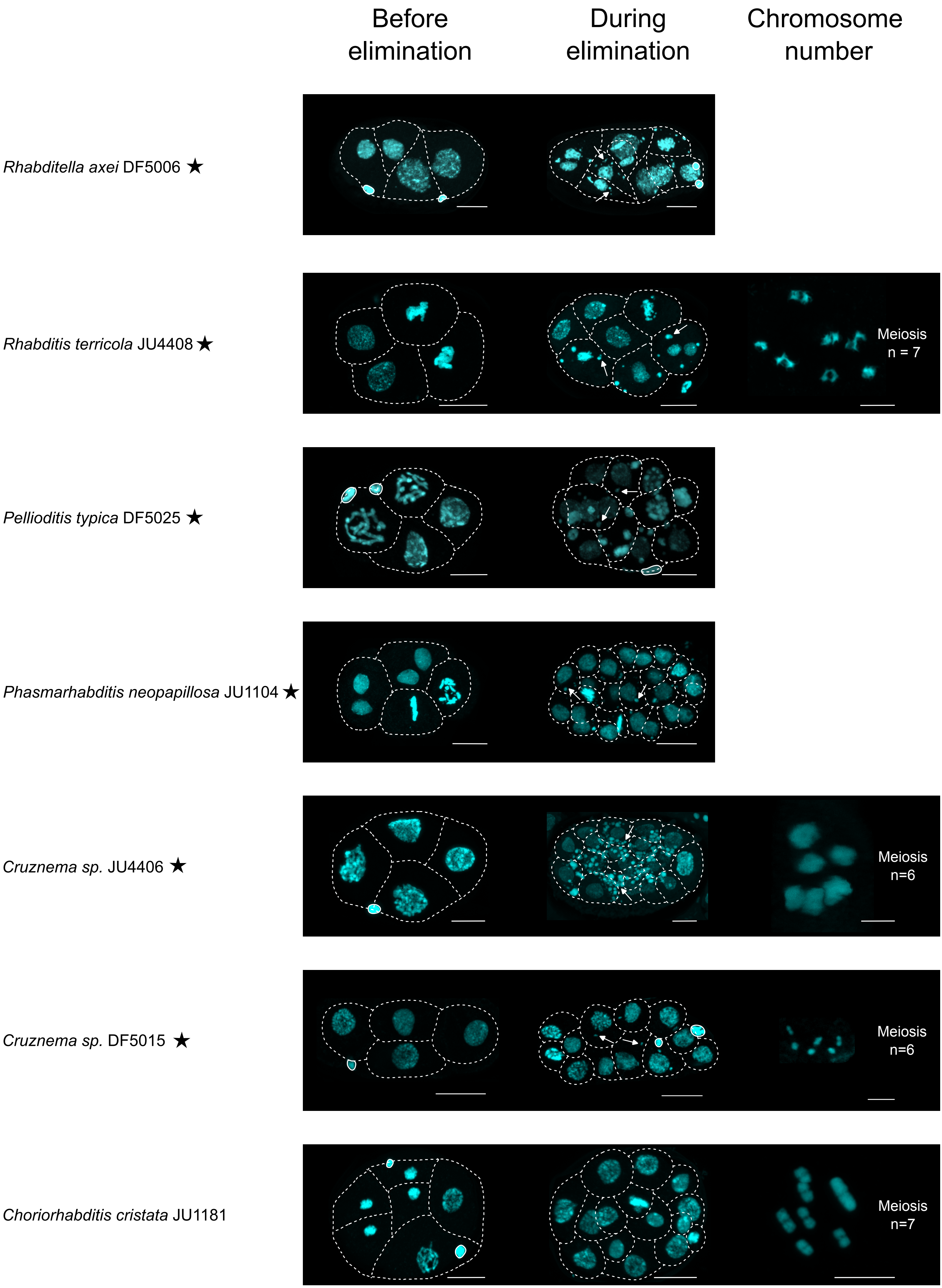

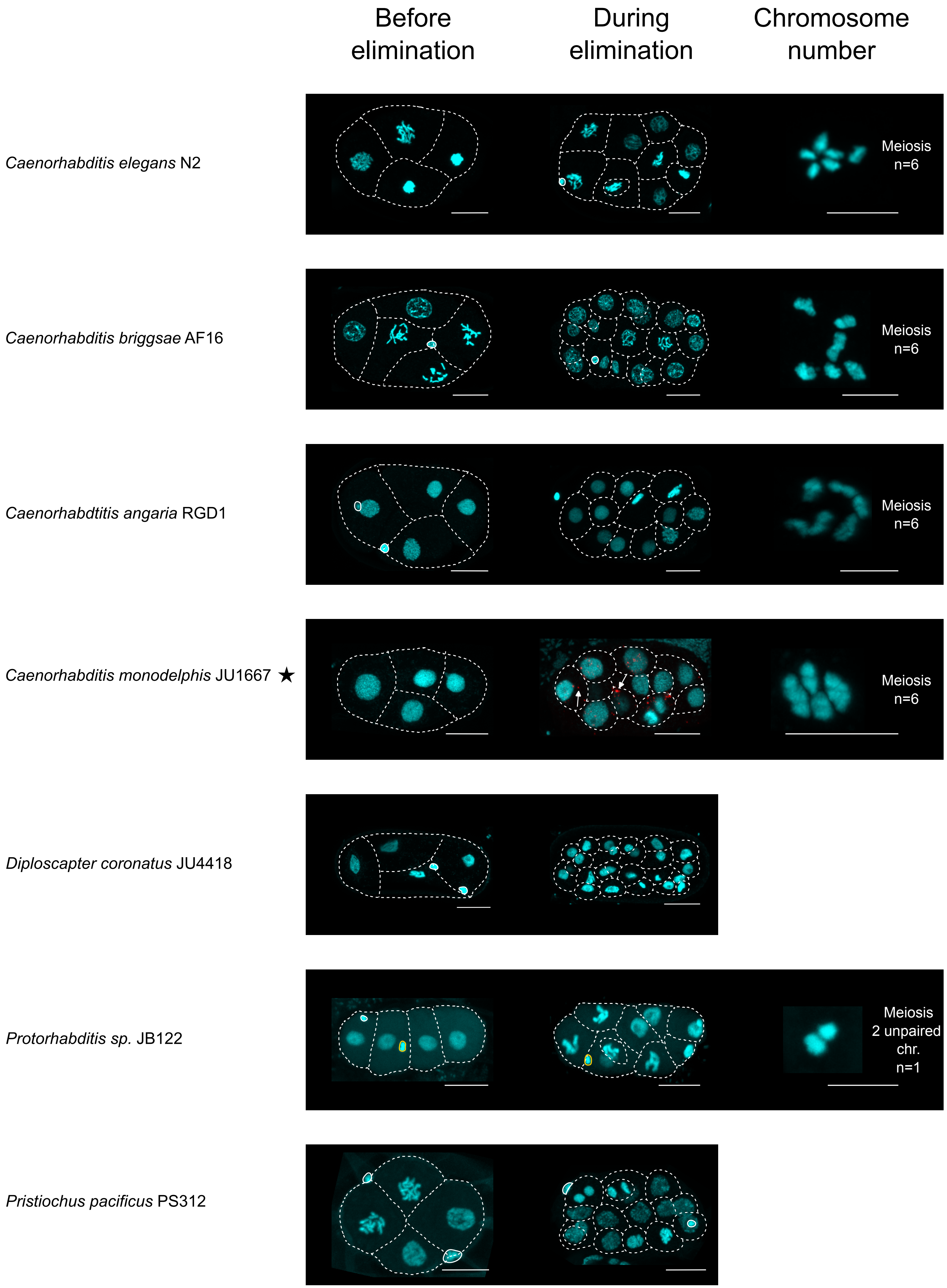

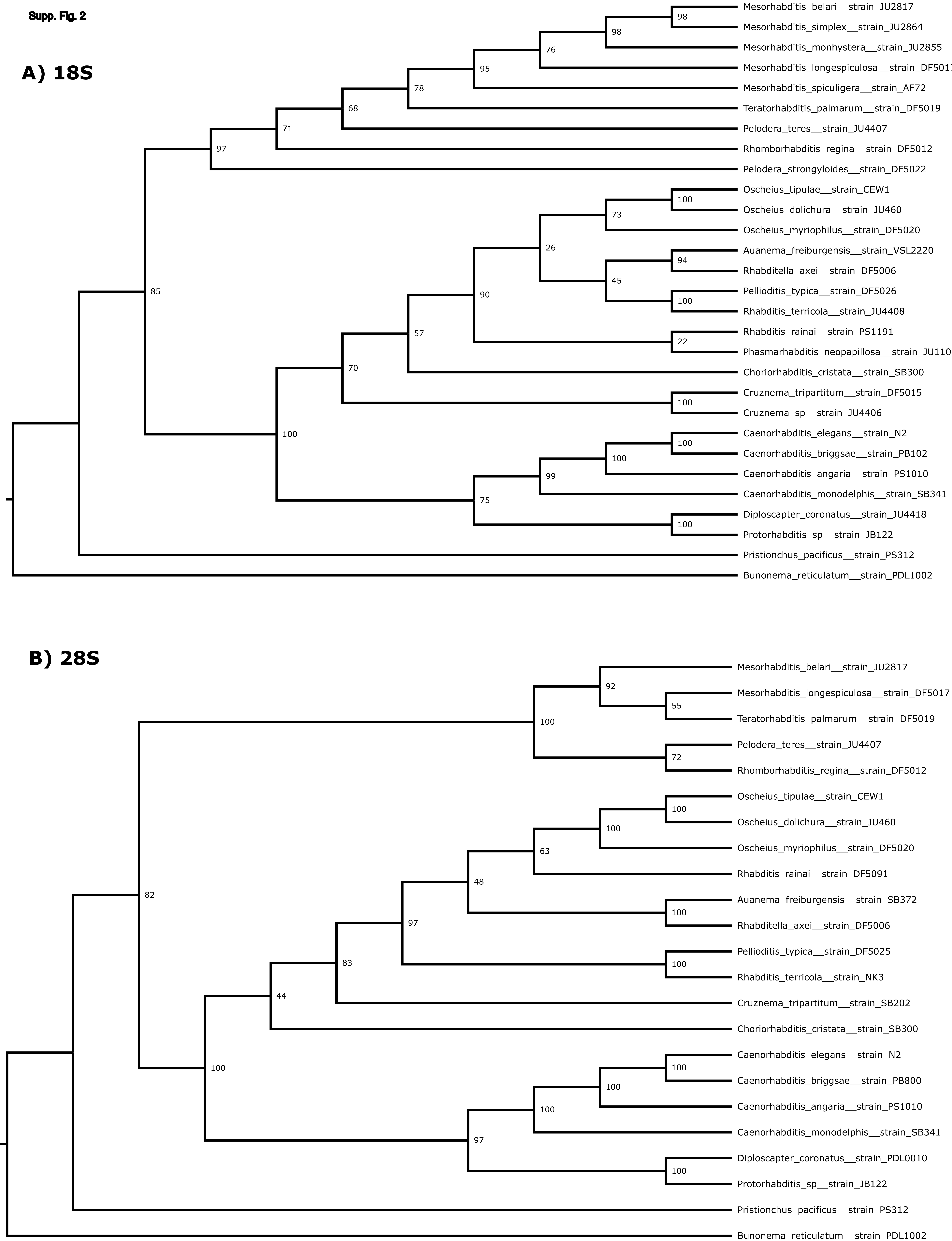
